# The interplay between dormant mutated cells and tumor promotion by chronic tissue damage in determining cancer risk

**DOI:** 10.1101/2024.01.24.577147

**Authors:** Yun Rose Li, Eve Kandyba, Kyle Halliwill, Reyno Delrosario, Quan Tran, Nora Bayani, Di Wu, Olga Mirzoeva, Melissa McCreery Reeves, Mishu Islam, Laura Riva, Eric Bergstrom, Kavya Achanta, John Digiovanni, Ludmil Alexandrov, Allan Balmain

## Abstract

While the causal role of mutagenic carcinogens in tumor development is well established, the relative contribution of environmental tumor promoting factors, wounding, and chronic inflammation is still unclear. Recent sequencing studies have suggested that most environmental carcinogens act as promoters rather than through mechanisms that involve direct induction of point mutations, but whether cancer risk factors such as obesity, chronic inflammation, wounding, or tumor promoters contribute directly or indirectly to mutation burden, or induce novel signatures, has not been investigated. Here, we present WGS analysis of over 100 mouse skin tumors to compare the effects of exposure to mutagens, the tumor promoter TPA, chronic wounding, obesity, or chemotherapy, on mutational burden and cancer risk. All tumors initiated by the carcinogen Dimethylbenzanthracene (DMBA) show a very strong A>T mutational signature (SBS.DMBA) attributable to a single exposure to this carcinogen. The number of SBS.DMBA mutations also showed a strong correlation with the “clock” signature SBS5, suggesting that one treatment with this mutagen can induce mutational signatures attributed to endogenous processes. No specific signatures could be attributed to obesity, high fat diet, wounding, or TPA. Cells carrying thousands of mutations persist over very long periods without inducing tumors or causing pathological changes but can give rise to tumors after short term exposure to TPA. Furthermore, normal cell turnover and proliferation during fetal and adult growth, is not sufficient for promotion, but tissue damage followed by regenerative proliferation seems to be required for tumor development. We conclude that tumor promoters, chronic inflammation, wounding, and obesity do not contribute significantly to tumor mutational burden, and that the rate-limiting determinant of tumor growth is exposure to a tumor promoter rather than the nature or number of genomic point mutations. These data are highly relevant to the recent demonstration of persistent oncogenic mutations in histologically normal human tissues during ageing.

## Introduction

Genetic changes observed in human cancer genomes are comprised of mutations caused by exposure to environmental carcinogens, or that result from spontaneous errors in DNA replication(1). Whole genome and exome sequencing of tens of thousands of human tumors have revealed mutational signatures that are the composite result of specific patterns of single base substitution (SBS) mutations resulting from exposure to environmental factors such as UV light (SBS7), cigarette smoking (SBS4), and aflatoxin (SBS24)(2–4). Large scale projects have been undertaken to investigate the mutagenic effects of hundreds of known or suspected human environmental carcinogens in bacteria(5), human cells in vitro(5), animal models(6), or by large scale sequencing of human tumors from high-risk regions(7,8) in an attempt to associate specific environmental exposures to cancer mutations. In this way, it may be possible to go beyond epidemiological associations and demonstrate a definitive causal link between specific environmental insults and cancer development.

Cancer risk is also heavily impacted by many complex and potentially modifiable, but poorly understood, risk factors such as obesity, diet, alcohol, hot drinks, wounding or chronic inflammation which are often collectively referred to as “lifestyle factors” (9,10) Obesity is a major potentially modifiable cancer risk factor second only to smoking and is likely to become the most important contributor to cancer risk globally in the near future. However, we presently do not know whether these factors contribute to cancer risk by directly causing specific mutational signatures, by acting indirectly to increase mutation rate through endogenous processes such as oxidative stress, or whether they act through non-mutagenic mechanisms to promote tumor outgrowth.

The relative contributions of known mutagenic processes and these other risk factors to the overall cancer burden in humans is now being re-evaluated because of two emerging findings. First, abundant mutations have been found by deep sequencing of DNA in multiple normal human tissues including skin(11) esophagus(12,13) colon(14), liver(15) endometrium(16) and lung(17) The number of cells carrying these mutations is very high in the skin and oesophagus, where as many as 70-80% of the cells have been clonally selected(11–13). Both driver gene mutations and the mutational signatures detected in these normal cells are like those found in tumors arising in the same tissues. This raises the question of why these apparently dormant mutated cells rarely progress to actively growing lesions (18). Secondly, whole genome sequencing (WGS) of both animal and human tumors suggests that many known carcinogens do not exert their effects through a mutagenic mechanism. Analysis of mutational signatures in mouse tumors induced by chronic exposure to known or suspected environmental carcinogens, showed that very few of these chemicals increased mutation burden or induced specific mutational signatures in tumors of the liver, lung, stomach, or kidney(6). Similarly, sequencing of hundreds of human oesophageal carcinomas from countries with varying incidence rates failed to reveal a mutational signature that may indicate the presence of a specific causative mutagen responsible for increased susceptibility(8). These findings raise the possibility that a fundamental step in cancer development is exposure to non-mutagenic environmental or lifestyle factors that stimulate clonal selection of pre-existing mutant cells lying dormant in normal tissues.

In the present study, we analysed the potential effects of a range of cancer risk factors including obesity, high fat diet, wounding, chronic inflammation, or treatment with a chemotherapeutic agent, on mutational burden and signatures using a well-established mouse initiation/promotion model of multistage skin carcinogenesis. This model involves exposure to a low dose of the mutagen DMBA, which is not in itself sufficient to result in tumorigenesis in wild-type animals. However, when DMBA is followed by repetitive treatment with an agent that causes tissue damage and inflammation, tumors inevitably develop in the exposed tissue(19). Using this model, we show that a single exposure of normal skin to a mutagen induces a highly variable number of carcinogen-specific mutations, which persist long term in the skin but are not sufficient for tumorigenesis. Subsequent exposure of the initiated tissue to chronic wounding or inflammation is the rate-limiting step for full tumor development, but whole genome sequence analysis of these tumor models showed that these exogenous factors do not significantly increase point mutation burden or induce novel mutational signatures.

## RESULTS

### Genomic mutational landscapes across a range of mouse skin tumor models

We tested the impact of a range of factors on mutational signatures by whole genome sequencing of 107 squamous papillomas or carcinomas from five different tumor models as shown in **(Figure 1)**. In four of these models, we used DMBA to initiate tumorigenesis by causing a specific A>T mutation at codon 61 of the *Hras* gene(20). In contrast to previous studies of long-term chronic carcinogen exposure(6), the application of a single dose of the carcinogen allows us to be confident of the timing of the initiating mutation. Any mutations induced by subsequent treatment of the animals with the tumor promoter TPA (model A), as a consequence of age of exposure (B), dietary factors or obesity (C), or chemotherapy (D), should not show the specific DMBA signature which is dependent on formation of bulky adducts with adenosine residues(21). In this way, we sought to distinguish carcinogen-induced initiating mutations with a specific mutational signature, from those that occur spontaneously or that result from a range of interventions that are known to increase cancer risk. We also carried out WGS on skin tumors induced in the complete absence of any chemical exposures, by activation of mutant *Hras* or *Kras* alleles using Cre-recombinase under the control of Lgr6 or Lgr5 stem cell promoters(22), followed by wounding and chronic inflammation (E).

**Figure 1:**
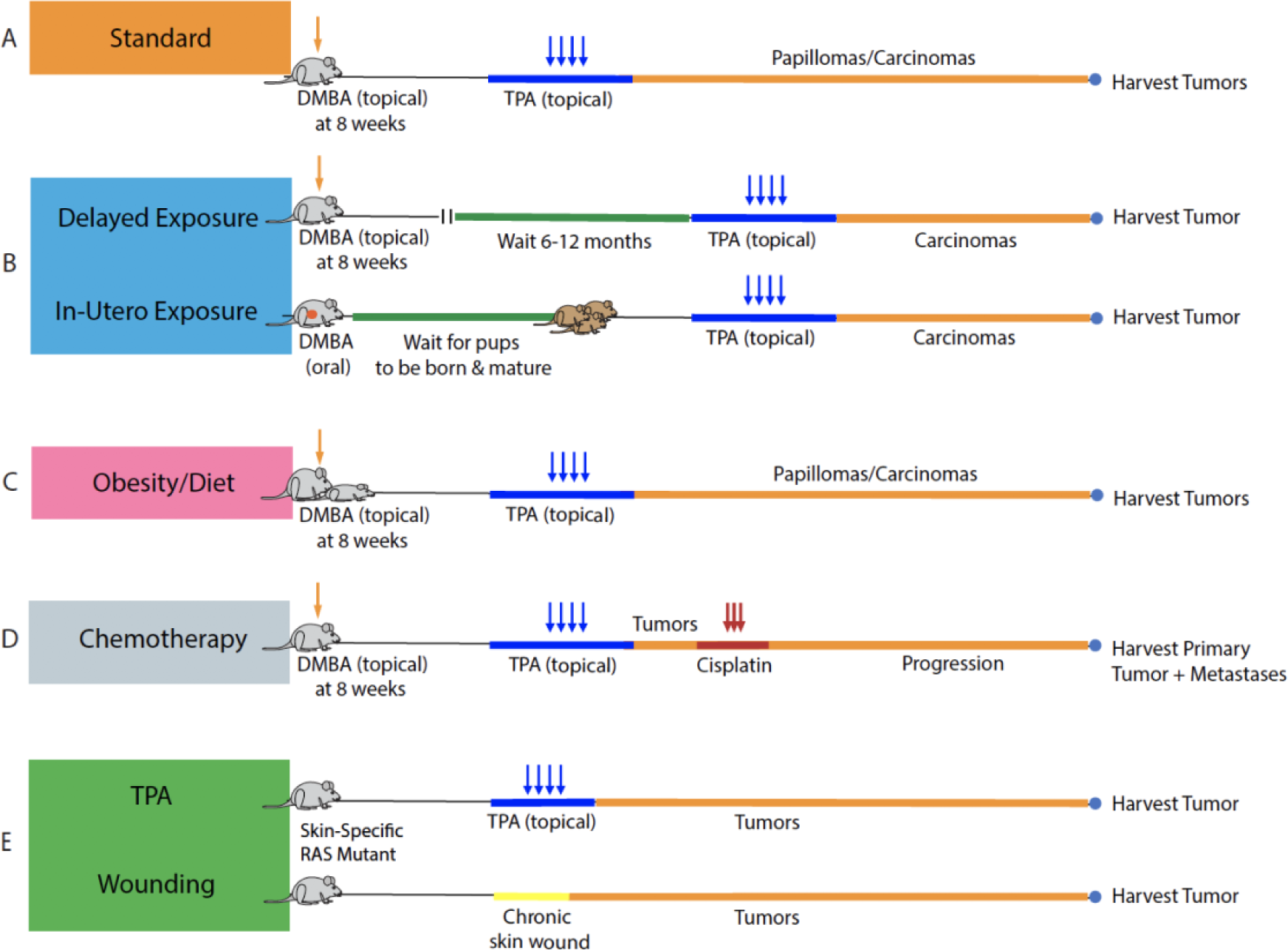
Mouse models involving carcinogen exposure together with a range of “lifestyle factors” important in human cancer aetiology. The majority of the mouse models utilized are based on the classical two-step skin carcinogenesis model as shown in (A) where adult WT mice are exposed to topical DMBA at 8 weeks of age followed by twice weekly administration of TPA for an additional 20 weeks. Mice are subsequently observed for tumor formation and sacrificed once tumor(s) are observed. In model (B) the original two step model is modified to investigate the consequence of delaying the administration of TPA by up to 6 months after DMBA exposure, and in a more extreme extension of this method, to investigate the consequence of administering DMBA orally to pregnant females resulting in “in-utero exposure” and subsequently administering TPA to the adult offspring (70). Model (C) depicts animals either genetically predisposed to obesity or leanness (*See Methods)* or WT animals exposed to either standard, calorically restricted, or high fat chow resulting in different BMI and consequent rates of tumor formation(33,34) when skin SCCs are induced using the same two-step model as depicted in (A). The Chemotherapy (D) model illustrates a model where WT animals with tumors induced by Model (A) are subsequently treated systemically by oral administration of cisplatin, mimicking human chemotherapy. In model (E) no mutagens are administered and animals with conditionally activated *Hras*/*Kras* driven by Lgr5/6 gene promoters are exposed to TPA promotion, or to chronic skin wounds. SCC (squamous cell carcinoma), DMBA (7,12-Dimethylbenz[a]anthracene), TPA (12-O-tetradecanoyl-phorbol-13-acetate), WT (wildtype), BMI (body mass index).

The total genome wide mutation burden seen in tumors from these models varies over 3 orders of magnitude (Range of SNVs from 20-48,425) **(Figure 2A, top panel)**. The lowest mutation burden is seen in skin tumors initiated genetically by the activation of *Hras* or *Kras* using Cre-recombinase under the control of the promoters of the stem-cell marker genes *Lgr5* or *Lgr6*(22). As previously demonstrated for genetically induced tumors of the lung(23), skin tumors initiated genetically had a very low overall mutation load (average 242.6 as compared to 6963 in DMBA induced tumors) **(Fig 2A and 2B, Sup Table 1)**.

**Figure 2:**
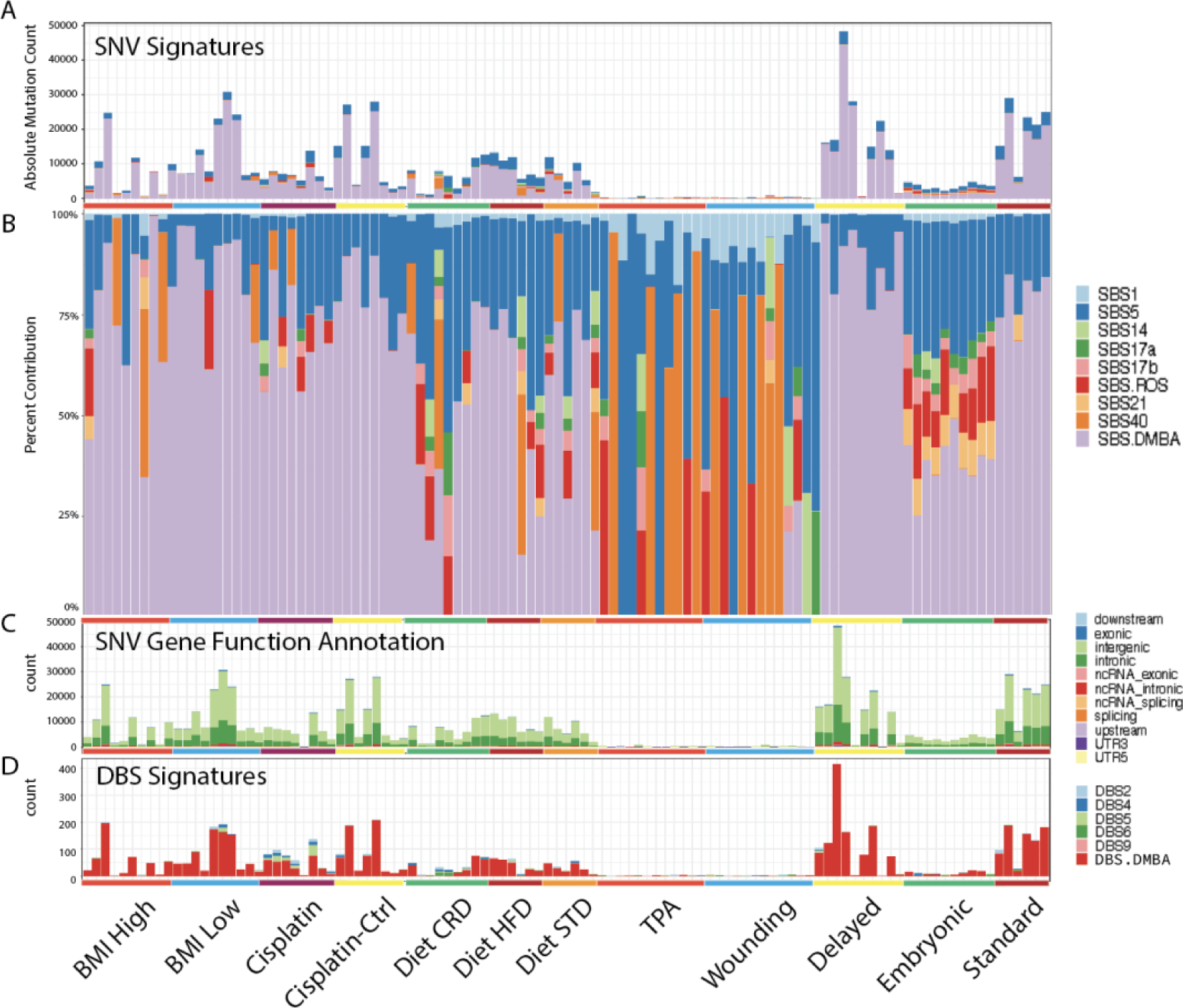
SNV mutations, functional mapping of mutations and mutational signature assignments using the COSMIC mutational signature database. A. Absolute SNV mutation load for each tumor (bars) with colours denoting assignment to each SBS mutational signature after decomposition to known mutational signatures in the COSMIC database. Note that the DMBA signature is a new signature not previously reported in the COSMIC human SBS database. B. Relative contribution of SNV mutations by COSMIC mutational signature type for each tumor sample. Note the prevalence of the SBS DMBA signature in nearly all samples with clear exception for the samples obtained from mice in the TPA and wounding promotion models. C. SNV mutational load for each tumor sample based on gene functional mapping to the mm10 genome. D. Double Base Substitution (DBS) signatures identified in the same samples including one attributable to DMBA (DBS-DMBA) and DBS5, attributed to Cisplatin exposure.

As compared with WGS analysis of 188 tumors induced by chronic exposure to 20 different environmental carcinogens, the total mutation burden in DMBA or genetically initiated tumors is more variable **(Sup Figure 1)**(6). The mutation load in the single low dose DMBA samples is comparable to or exceeds the total mutation burden induced by chronic exposure to the most highly mutagenic environmental carcinogens tested, for example TCP or vinylidene chloride. However, some DMBA samples had a very low mutation burden comparable to tumors induced by most other agents for which no mutation signatures were detected.

### Mutational Signatures in skin tumor models

We next carried out a detailed analysis of the mutational signatures found in skin tumors from the various models **(Figure 1 and 2)** using *SigProfiler* (See Methods)(24). In total, we extracted four *de novo* signatures, which decomposed to 9 known human COSMIC SBS mutational signatures(24) in addition to a mutational signature for DMBA **(Figure 3A-C)**. The dominant de novo mutational signature in all samples initiated using DMBA was initially decomposed into a combination of human SBS22 and SBS25. The former is associated with aristolochic acid exposure in several human solid tumor types(25). Both DMBA and aristolochic acid form adducts with adenosine residues in DNA, resulting in A>T transversion mutations(25,26).

**Figure 3.**
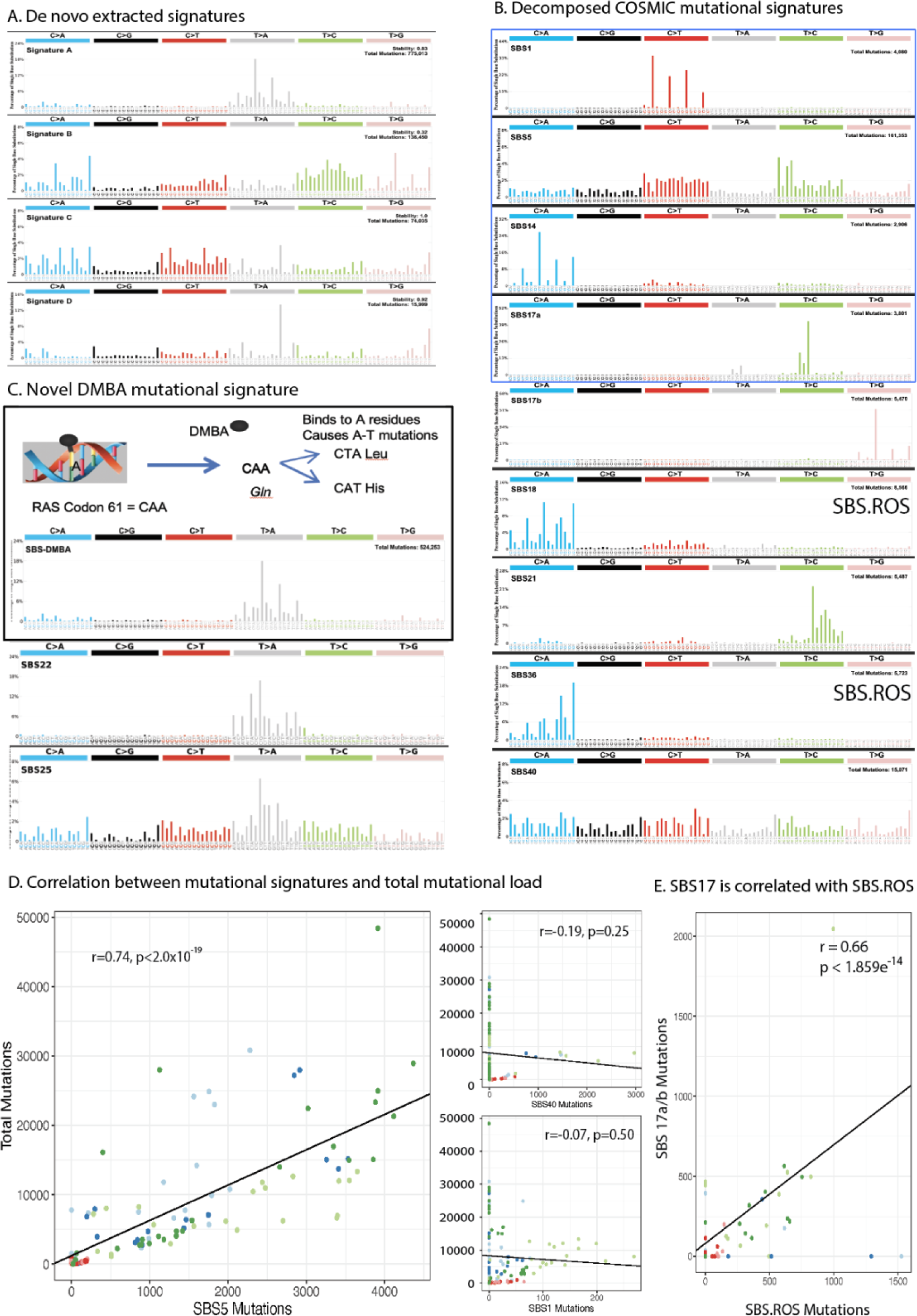
SNV Mutational Signatures Identified with SigProfiler. A. De novo SBS mutational signatures (4) identified based on WGS data from the entire tumor cohort. B. Molecular basis of DMBA induced mutational changes and comparison of novel SBS DMBA signature as compared to SBS22 and SBS25, two known COSMIC signatures that bare strong resemblance to the DMBA signature. C. Analysis of de novo SNV mutational signatures resulted in the decomposition to 9 known SBS COSMIC signatures plus the novel SBS DMBA signature. D. Strong correlation between mutations assigned to SBS5 and total mutation load across all tumor samples (Pearson’s r=0.74, p<2.0×10^−19^) while SBS 1 and SBS40 showed no such correlation. E. Correlation between frequency of observed mutations attributed to SBS.ROS versus SBS 17a and SBS 17b (r=0.66, p<1.86×10^−14^)

To identify specific differences between the DMBA signature and SBS22/25 we examined the pentanucleotide sequence rather than the trinucleotide sequence context of these mutations (*See methods*). This analysis showed that these signatures can be distinguished at the pentanucleotide level, and we therefore identified SBS.DMBA as a specific signature of exposure to DMBA (Fig 3C). This mutational signature identifies mutations induced at the specific time point of DMBA exposure, in contrast to mutations attributed to other endogenous processes such as proliferation or inflammation, which are much more likely to occur during subsequent tumor promotion or progression. Interestingly, this analysis also detected a double base substitution signature (DBS-DMBA) that was also only detected in samples initiated using this carcinogen **(Figure 2D and Sup Fig 3)**.

Several other mutational signatures were identified in mouse tumors, including the clock-like signatures SBS1,40, and 5, as well as additional signatures that have been attributed to the activity of reactive oxygen species or chronic inflammation (SBS18, 36, 17a and 17)(2,27). However, no additional novel mouse-specific signatures were identified that could be attributed to the interventions shown in Figure 1. To investigate the possibility that some mutational signatures could be indirectly caused by DMBA, for example as a response to stress induced by carcinogen exposure, we carried out an analysis of the relationship between the total number of DMBA-induced mutations in each sample, and the numbers of mutations associated with other signatures. Surprisingly, this analysis showed a very strong correlation between the total number of SBS.DMBA mutations and those attributed to the “clock-like” signature SBS5 **(Figure 3D, Spearman’s r=0.74, p<2.0e-19)**. However, the same observation does not extend to the clock signatures SBS1 and SBS40 (spearman’s rho =0.07 and 0.19, respectively).

Since SBS5 is almost ubiquitous in human cancers, this association cannot be specifically due to the mis-repair of DNA adducts induced by the carcinogen DMBA but may be a consequence of more generalized genomic instability that is proportional to the total levels of damage induced by DMBA, resulting in activation of fragile sites in the genome. Notably, carcinogen-induced DNA damage can lead to genomic changes or aberrant transcripts in the fragile site gene *Fhit,* and germline deletion of this gene in the mouse leads to development of tumors with high levels of the SBS5 mutational signature(28). An association between total DNA damage, as monitored by mutational burden from WGS analysis, and the levels of SBS5 was also seen in human tumors from TCGA, and in mouse tumors induced by chronic exposure to a range of carcinogenic chemicals **(Sup Table 2)**(6) We conclude that in addition to a specific mutational signature attributable to DMBA-DNA adduct formation and mis-repair, a more general DNA damage response may result in additional mutations with the SBS5 mutational signature.

### Analysis of signatures due to induction of ROS

An important endogenous source of mutations in human tumor genomes is reactive oxygen species (ROS), a broad category of highly reactive oxygen free radicals including peroxides, superoxide, hydroxyl radical, singlet oxygen, and alpha-oxygen(29). ROS signatures SBS18 and 36 show strong cosine similarity and both demonstrate strong predominance of G to T transversions. We noted that on iterative runs of analysis depending on which samples are included, de novo observed mutational signatures can be decomposed to either SBS18 or 36, and thus appear to be computationally interchangeable. We therefore designated mutations assigned to either SBS18 or SBS36 as SBS.ROS. Correlation analysis across all samples for ROS signatures 17a, 17b, 18, and 36 shows that, as expected, SBS17a and SBS17 b are strongly correlated (r=0.87, p<1.33e-33), but more importantly, we identified a strong correlation between the sum of mutations assigned to SBS17a and SBS18b with sum of mutations assigned to SBS.ROS (spearman’s correlation r=0.66, p<1.86e-14) **(Figure 3E and Sup Figure 5)**. This suggests that the induction of DNA damage by ROS stimulates several independent processes leading to DNA repair and residual DNA changes, resulting in induction of distinct ROS-associated mutational signatures.

**Figures 2A and B** show that SBS.ROS, as well as SBS17a and 17b, were present at variable levels in many samples investigated. In the embryonic treatment cohort, almost every tumor sample exhibited a significant mutation contribution from SBS.ROS, 17a and/or 17b **(Sup Figure 5B)**. This seems likely to be due to the fact that in this cohort of mice, pregnant females were treated systemically with DMBA at a stage of development when the skin of the pups is highly proliferative. It is possible that induction of a DNA damage response in skin during active proliferation, in contrast to 8-week-old adult mouse skin which is largely quiescent, results in a greater degree of oxidative damage. Further support for this possibility came from analysis of mutational signatures in tumors from mice that were treated with cisplatin. Carcinomas from mice that had been treated at age 15 weeks with 3 doses of cisplatin showed a clear increase in SBS.ROS and SBS17a/b mutations, together with the known human cisplatin-associated dinucleotide mutational signature (DBS5) **(Figure 4C and D and Sup Figure 3)**. This observation is compatible with the interpretation that actively proliferating normal, or tumor cells may be more susceptible to ROS mediated mutagenesis upon exposure to a DNA-damaging factor, but further studies would be required to confirm this association.

**Figure 4.**
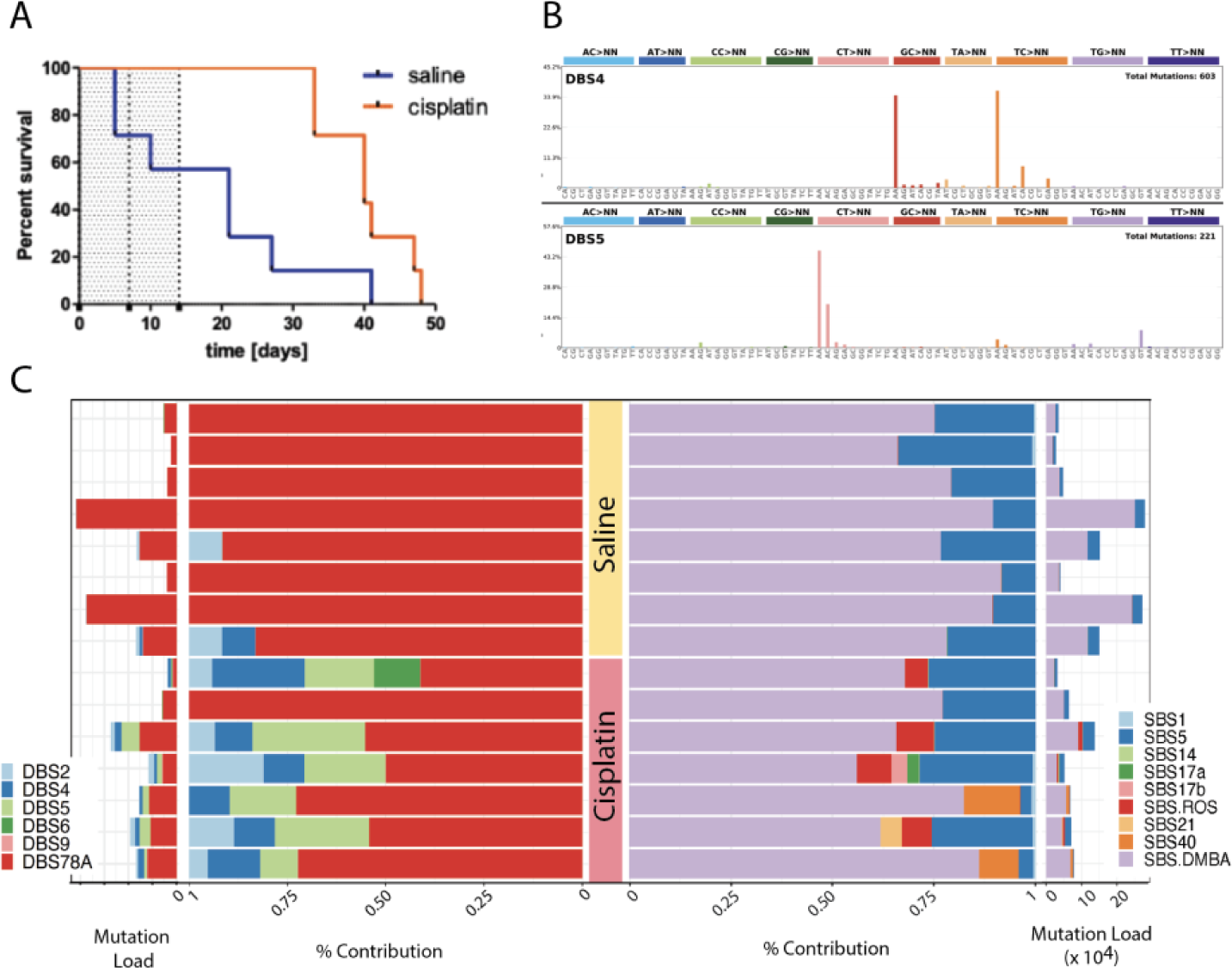
Cisplatin induces human DBS signature and SBS.ROS. A. Tumor free survival difference between mice with tumors induced by the two-step model either treated with cisplatin (Orange) or treated with saline (Blue). See Figure 1 for experimental design. B. Two DBS mutation patterns identified in cisplatin treated tumors but not in saline treated tumors. SBS18 and 36 are found in cisplatin treated tumous but not in saline treated samples. C. DBS (Left) and SBS (Right) mutation signature analysis results. Each side shows both total mutational load (thin panel) and relative contribution of each signature to total mutational load

Finally, we considered the possibility that a high DMBA mutation load could potentially confound the detection of ROS signatures, particularly as DMBA may also induce G>T mutations that mask SBS18 and SBS36. We repeated the signature extraction after removal of 90% of T>A mutations scaled by the relative trinucleotide frequencies observed in the DMBA signature (*see Methods*). We found that the high relative contribution of SBS.ROS mutations in tumors from embryonic cohort mice was not affected by removal of DMBA mutations, although SBS.ROS mutations were more consistently detected in the adult DMBA/TPA treatment groups using this method **(Sup Figure 2)**. We did not observe a clear correlation correlation between any of the insertion-deletion signatures identified with DMBA exposure **(Sup Figure 4)**.

### Genetically or chemically initiated cells persist long term in mouse skin and require a promoting stimulus for tumor development

Mice in which *Hras* or *Kras* mutations were initiated genetically rather than by chemical mutagens **(Figure 1, model E)** also developed aggressive tumors, but only after chronic exposure to TPA, or in response to full thickness skin wounding **(Sup Figure 8A)**. The combination of activated *Kras* and wounding was particularly potent, resulting in large skin tumors, predominantly at the sites where wounds were clipped, that required the mice to be euthanised after 6-8 weeks. No obvious differences were found by WGS analysis of tumors resulting from *Hras* or *Kras* mutations, or that were related to the cell of origin (*Lgr5* or *Lgr6*-positive stem cells) **(Figure 2B and Sup Figure 8B)**. No tumors developed in these animals by activation of *Ras* mutations with no further treatment for at least 5-6 months, as previously shown after delivery of mutant *Ras* by retroviral oncogene delivery to the back skin of mice(30). Notably, genetically initiated tumors did not carry any novel mutational signatures that could be attributed to the activity of either TPA or wounding, and all carried the endogenous signatures SBS1, 5, and/or 40, with variable contributions from the ROS signatures SBS.ROS, 17a or 17b **(Figure 2B, Sup Figure 8)**. Very few large-scale translocations were observed in these samples as well **(Sup Figure 9)**.

Analysis of mutational burden and signatures in tumors induced using model B, in which initiation was followed by a long delay before starting tumor promotion using TPA, demonstrated that initiated cells persist long term in the skin without causing any obvious pathology, as previously suggested by early studies on the rate-limiting role of promotion in multistage carcinogenesis(31). Importantly, our data show that DMBA-initiated cells carrying thousands of DMBA-signature mutations, as well as those carrying a single genetically induced *Ras* mutation, were persistent in the skin, but remain sensitive to the effects of tumor promoters.

### Obesity models do not show new mutational signatures, but gene expression changes are associated with high/low BMI

We performed WGS of SCCs associated with genetically determined high or low body mass index(32) or diet-induced obesity(33,34). We have previously shown that interspecific backcross mice ((SPRET/Ei X FVB) x FVB) show a wide distribution in body weight and weight/length ratio (henceforth high and low BMI groups; **(Figure 5A)**. Mice of the high BMI group, in particular, male mice of this cohort, develop earlier onset and a higher incidence of skin tumors compared to low BMI mice of the same backcross population(32) **(Figure 5B)**. To complement this model of genetically induced high or low BMI, we also investigated a model of environmentally induced obesity, by sequencing tumors induced in mice that were fed either standard, high fat, or calorically restricted diet(33,34). While we detected a wide range of mutation burdens in these tumors, no significant differences in mutation load or mutation signatures were observed, although there was a trend for higher mutation loads in the tumors from the low compared to high BMI group (Student’s T-test, p=0.11).

**Figure 5.**
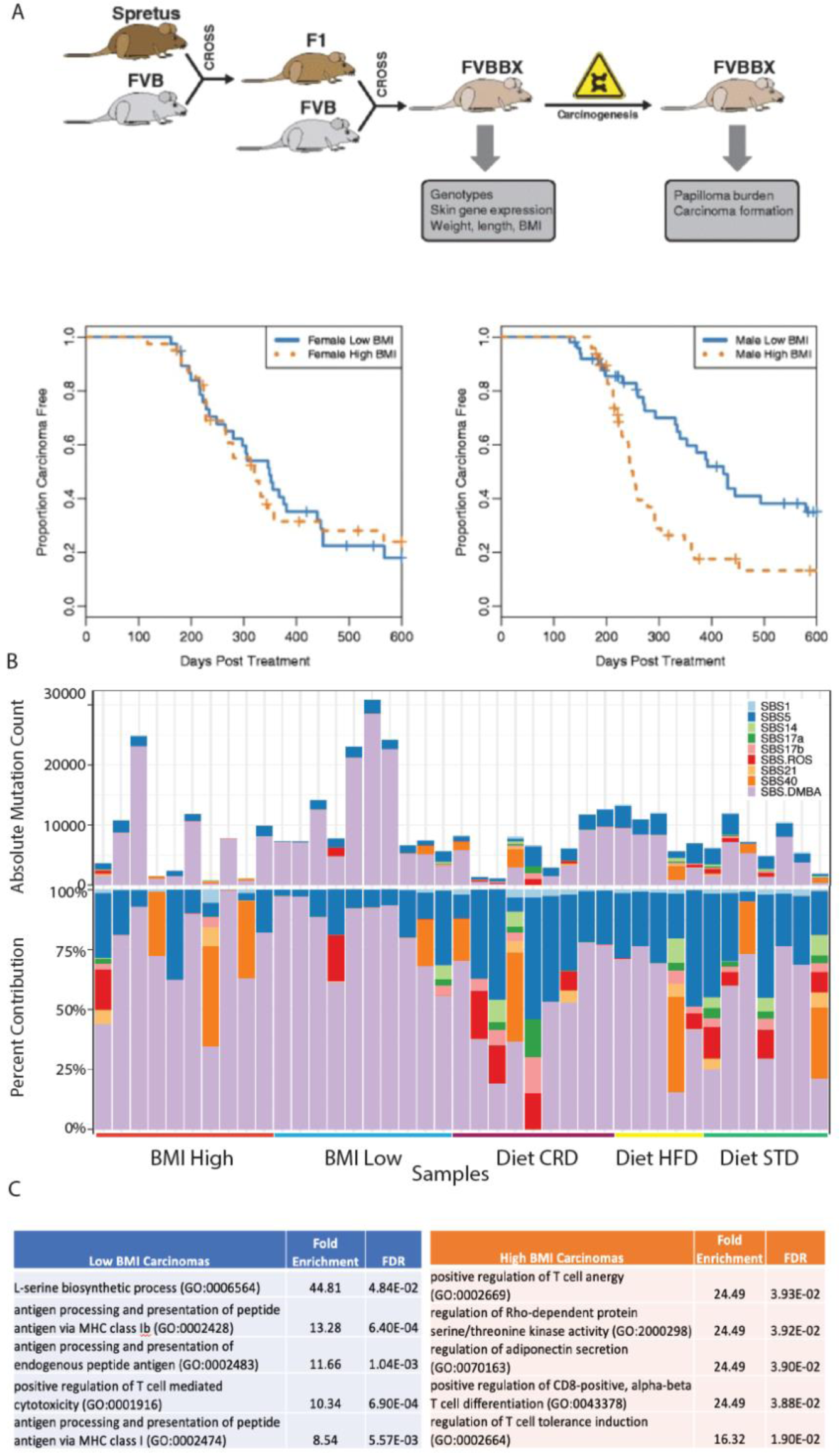
Genetic or dietary obesity impacts gene expression, but not mutation burden or mutational signatures. A. Schematic describing the genetically derived high versus low BMI animals (ABOVE) and the sexually dimorphic differences in rate of tumor development in high BMI males but not females (BELOW) B. Absolute SNV mutations mapping to COSMIC SBS mutational signatures including those attributed to SBS DMBA (ABOVE) and relative contribution of mutations assigned to COSMIC signatures excluding those attributed to SBS DMBA (BELOW). C. Significant biological pathways identified by pathway analysis of the most strongly enriched genes by gene expression profiling in male mice (Genes highly expressed in high BMI mice SCC and Genes highly expressed in low BMI SCC)

Since no significant differences were observed in total mutation burden or mutational signatures attributed to DMBA or ROS amongst animals in either the genetically determined or dietary obesity cohorts, we speculated that obesity or a high fat diet may cause changes in gene expression resulting in increased tumor growth capacity. To test this possibility, we carried out microarray analysis of gene expression in squamous cell skin tumors in the high (n=36) vs low (n=41) BMI groups. We found significant enrichment for pathways such as antigen processing and MHC peptide antigen presentation and positive regulation of T cell mediated cytotoxicity in Low BMI carcinomas. In contrast, in the high BMI tumors we observed enriched expression for genes in the T-cell anergy pathway and regulation of T cell tolerance induction. Future single cell transcriptome studies would be necessary to tease apart whether the expression identified here is due to intrinsic tumor cell expression or tumor infiltrating immune cells.

### Driver gene mutations in skin tumor models

We observed a broad range of mutations that mapped to driver genes in mouse SCCs from this cohort which have been previously reported as recurrently mutated in human cancers(35,36) **(Figure 6 and Sup Figure 6)**. To examine the contribution of specific mutational processes to driver mutations, we examined the probability that a given driver mutation could be attributed to a specific mutational signature **(Figure 6B)** (*See Methods*). We observed that driver mutations in a variety of genes from chemically induced tumors have incurred DMBA-associated A>T mutations at the same time as *Hras* during initiation; notably, no such driver mutation in previously reported human oncogenes were detected in tumor samples that were not initiated by DMBA **(Sup Figure 8)**. The A>T mutations were seen in genes encoding membrane proteins Fat1-4, tumor suppressors involved in cell-cell communication and Hippo/Yap signalling (37) as well as Card11, involved in Nfkb signalling and immune responses(38). These genes, as well as many of the other genes in **(Figure 6)**, including epigenetic regulators Kmt2a-d, are commonly mutated in human squamous tumors of the skin, oesophagus or head and neck(39,40).

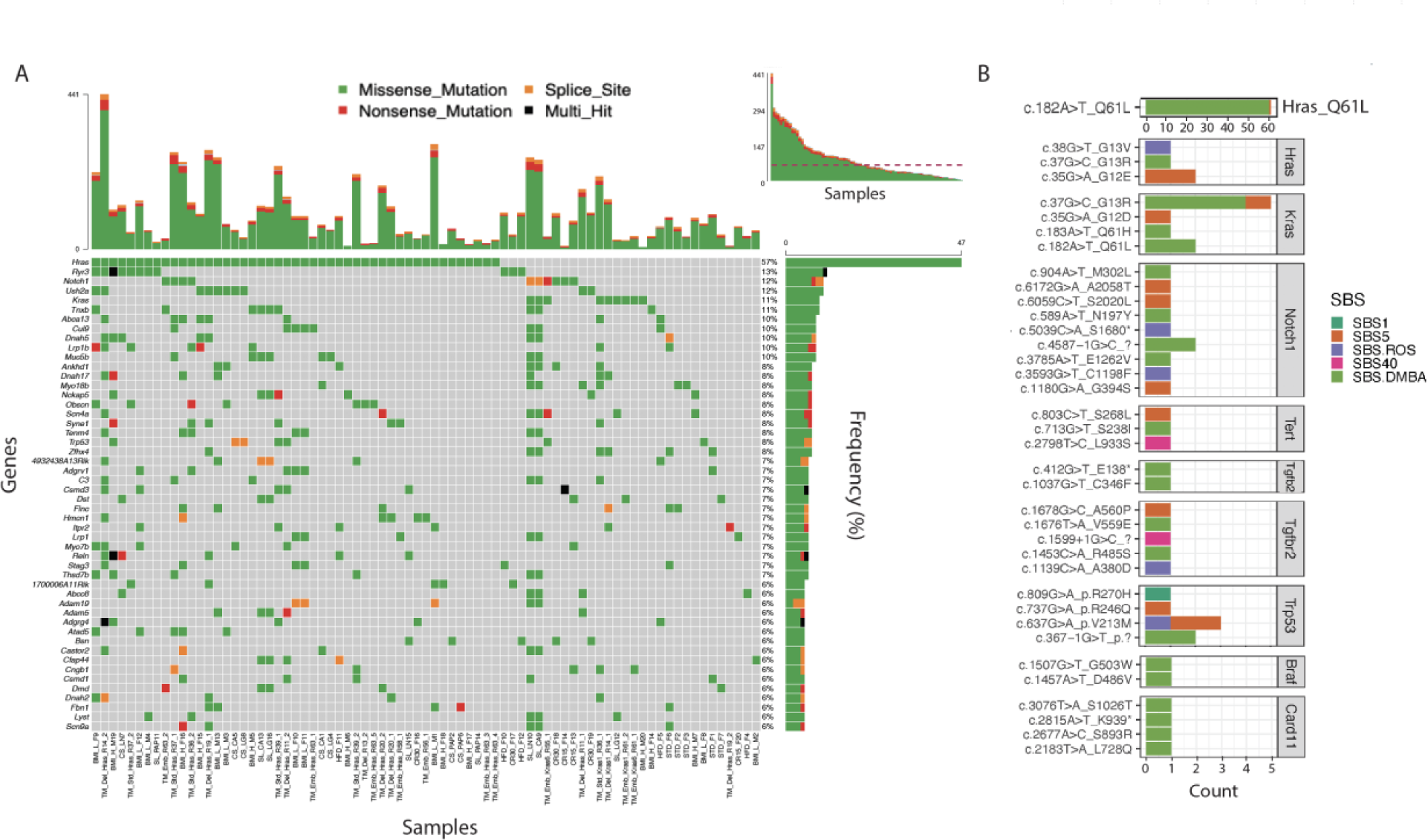

Although most recurrent driver mutations display the DMBA mutational signature, some tumors had driver mutations that were associated with other detected signatures including SBS5, SBS1, SBS18 or SBS36. Some of these mutations may be attributed to endogenous processes activated at the time of initiation (e.g. SBS5-see discussion above) or alternatively by ROS generation, leading to mutations with SBS18 or 36 signatures during the promotion or progression phases of carcinogenesis. Gene mutations associated with these other processes were found in *Trp53, Notch1, Tert*, and *Tgfbr2*, all of which are also commonly mutated in human squamous tumors(41).

We next compared the driver mutations observed in the initiation/promotion models of carcinogenesis in the skin to the solid tumors induced in different tissues by chronic low dose exposure to suspected human carcinogens we previously examined(6). In contrast to the uniform patterns of initiating mutations seen in the two-step DMBA/TPA model, which were clearly attributable to the mutational signature induced by DMBA, tumors induced in the forestomach by Trichloropropane (TCP) or in the kidney by Vinylidene Chloride (VC), did not have a high frequency of driver mutations specifically attributable to the mutational signatures directly caused by exposure to these agents(6). Closer inspection of these data however showed that although the specific mutations were not obviously attributable to the direct mutagenic action of each carcinogen, there was clearly carcinogen-specific selection of driver mutations by different exposures to the various non-mutagenic carcinogens (**Sup Figure 7**, see also Discussion).

## DISCUSSION

We have demonstrated here that the number of somatic mutations induced in normal mouse skin by a single treatment with the mutagen DMBA, either during embryonic development or in adults, varies over 2-3 orders of magnitude, but is clearly insufficient for tumor formation in the absence of a promoting stimulus of tissue damage by TPA. This situation mirrors the observations of abundant mutations in normal human tissues that do not cause any obvious pathological changes(11–13). In the mouse, cells carrying thousands of SBS.DMBA signature mutations persist over long periods of time in normal tissue, but only give rise to actively growing lesions after exposure to 1-2 months of treatment(42) with the promoting agent **(Figure 1,2)**. DMBA-initiated cells in normal skin clearly have enough mutations, and in the correct combinations, for eventual tumor development, but have an absolute requirement for the promotion stage.

Our analysis of whole genome sequences of 107 squamous tumors initiated by activation of mutant Ras, indicate that the contribution of promotion processes, including treatment with TPA, wounding and chronic inflammation, obesity or a high fat diet, to total mutation burden is minimal. Normal cell proliferation, for example as seen during fetal development, neonatal growth and adult homeostasis, does not seem to be sufficient for completion of the promotion phase, as mice exposed to DMBA initiation *in utero* do not develop papillomas even after the large number of cell divisions required for foetal and adult growth. Our data support the conclusion that promotion requires a regenerative type of cell division, such as that seen in the adult skin during wound healing or growth in response to repetitive tissue damage by TPA.

Although the majority of point mutations found by WGS analysis of skin tumors initiated with DMBA are attributable to the direct action of the initiating mutagen DMBA, some rare mutations were found in established driver genes such as the *Trp53* tumor suppressor gene, or in Transforming growth factor beta receptor 2 (*Tgfbr2*) that did not carry the SBS-DMBA signature but rather had mutations more likely attributable to clock-like signatures or SBS.ROS **(Figure 6)**. *Trp53* mutations are known to occur in late stages of tumor development in this model(43), and TGFB signalling is also important for late-stage tumor invasion(44). Notably, all of the tumors that had mutations in *Tgfbr2* were from mice subjected to caloric restriction, or that had a very low body mass index in the genetic BMI model **(Figure 1C)**. TGFB signaling plays an important role in insulin signaling(45), thus it is possible that metabolic restriction may have resulted in selection of cells carrying mutations in this pathway(46). ROS signatures were more consistently detected in tumors arising from models, for example the in-utero exposure model B, or the cisplatin chemotherapy model D **(Figure 1)** that involved exposure of proliferating cells to DMBA, suggesting that DNA damage during active cell proliferation may be more likely to induce reactive oxygen species. We conclude that while the overall contribution of promotional processes to mutation burden is very low, some of the rare mutations that arise during promotion may be positively selected and could facilitate later stages of malignant progression. The lack of a major contribution to SNV mutational burden does not preclude an important role for other forms of DNA damage, for example induction of aneuploidy or gross chromosome changes, that could be induced by tissue damage or chemical tumor promoters.

In fact, early studies of the effects of TPA on cells in culture identified a “clastogenic” effect of TPA treatment, resulting in chromosome aberrations (47,48). *In vivo*, chronic TPA treatment resulted in appearance of papillomas showing aneuploidy of chromosomes 7 and 6 in pre-malignant papilloma cells(49) and tumors induced by DMBA/TPA rather than repeated DMBA treatment showed a higher level of chromosomal alterations(50). Further studies of links between tumor promoters, wound healing processes, and gross chromosome instability will be required to address these questions.

Genetic activation of mutant *Kras* or *Hras* expression in the skin did not lead to tumor formation over several months, but a single full thickness wound leads to induction of rapidly growing lesions within a few weeks. These tumors almost completely lacked point mutations in known cancer drivers other than the engineered mutations in Ras genes, as previously noted for other engineered mouse cancer models(23). It is possible that direct induction of mutations in Lgr5+ or Lgr6+ stem cells may abrogate the requirement for many other mutations, or alternatively that simultaneous activation of *Ras* mutations in multiple adjacent cells in the same compartment may allow initiated cells to escape competitive inhibition by surrounding normal cells(51). The strong selection of Ras mutated cells after a wounding stimulus is in agreement with historical chemical carcinogenesis studies (52), as well as with other models demonstrating clonal selection of cells expressing mutant Ras genes by tissue damage or tumor promoter treatment (53)(54)(55).These results however differ from other models targeting mouse ear skin, in which normal cells can out-compete their mutant counterparts after wounding (56). These differences may be due to the differing experimental design resulting in altered balance between numbers of normal and mutant cells, altering the competitive landscape to favor the selection of normal cells.

Our data, together with results of previous work on WGS of mouse tumor models(6,18) emphasize the potential importance of environmental promoting factors as agents of selection of pre-existing cells carrying driver mutations. Examples of promoter-induced selection of different driver mutations have previously been reported for skin (57) and liver tumors(58). In the previous study by Riva et al(6), spontaneously arising tumors had the highest frequency of Braf V637E (equivalent to human V600E) mutations, but this driver mutation is very rarely found in tumors promoted by a variety of different non-mutagenic promoting factors (p<0.012, Fisher’s exact test). Another common spontaneous liver mutation was *HrasG13R* (in 5/12 spontaneous tumors) but tumors induced by exogenous agents only rarely (4.9%) had the *HrasG13R* mutation, with most displaying mutations in alternative locations in *Hras* or *Ctnnb1* **(Sup Figure 7)**. In the lung, only 3 of 14 spontaneous tumors had mutations in *Kras*, whereas 59.5% of all carcinogen-promoted lung tumors showed mutations in this strong cancer driver gene. We conclude that different types of exogenous factors damage normal tissues in distinct ways and stimulate tissue regeneration through alternative pathways that select from a wide array of pre-existing spontaneous or carcinogen-induced mutations. The recent observations of abundant potential driver mutations in completely normal human tissues supports the hypothesis that environmental promoting factors play a major role in human cancer aetiology. Strong support for this possibility was recently published by Hill et al (59), who demonstrated that air pollution, which is known to be associated with lung cancer in non-smokers, induces tumors with a low mutational burden and frequent mutations in the *EGFR* cancer driver gene. Further studies from this laboratory(60) have identified long-lived stem cells expressing *Lgr6* in the upper hair follicle of mouse skin, rather than classical bulge stem cells, as major cells of origin of chemically induced skin tumors that can remain dormant over long periods, but still retain the capacity to form tumors after exposure to tumor promoters. These results confirm the observations by Berenblum and Shubik (31,61) that initiated cells can remain in the skin permanently, and that the most important and rate-limiting step in carcinogenesis was repeated exposure to promoting factors rather than initiation.

## Online Methods

### Mouse breeding, husbandry and tumor induction

#### A. BMI Backcross Model (Genetic Model)

This experiment has been described previously in detail in Halliwell et al(32). Briefly, male *Mus spretus* (SPRET/Ei) and female *Mus musculus* (FVB/N) mice obtained from the Jackson Laboratory were crossed to generate F1 hybrids. Female F1 mice were then crossed to male FVB/N mice to generate backcrossed mice, referred to as FVBBX mice. Tail tips were taken at seven weeks and snap frozen in liquid nitrogen for RNA and DNA. For mice not undergoing tumor induction, tail, dorsal skin, and other tissues were harvested after sacrifice by asphyxiation and cervical dislocation at seven weeks.

Chemical carcinogenesis was initiated using the two-stage DMBA/TPA protocol. In brief, seven-week-old mice were shaved and treated with 25 μg DMBA in acetone over an approximately 2-in. squared region on the center of the back. For the following 20 weeks, 200 μL of 10−4 M TPA in acetone was applied twice per week to the DMBA-treated area. Mice were shaved two days prior to the first TPA treatment for that week. TPA treatments were continued for 20 weeks.

#### B. High fat diet/caloric restriction (Dietary Model)

This experiment has been described previously in detail(62). Briefly ICR female mice (3–4 weeks of age, Harlan Teklad) were placed on the 10 kcal% fat control diet at 7 weeks of age and initiated with 25 nmol of 7,12-dimethylbenz[a]anthracene (DMBA; Eastman Kodak Co.). Four weeks following initiation, mice were randomized and placed on the 4 diets (n = 30 per group). The three groups consisted of control (10 kcal% fat control diet), high fat diet (70 kcal% high fat diet), and caloric restriction (15 or 30% caloric restriction). Four weeks later, mice received twice weekly topical treatments of 3.4 nmol of TPA (Alexis Biochemicals) for 50 weeks. Mice were weighed before randomization and then every 2 weeks for the duration of the experiments. Tumor incidence (percentage of mice with papillomas) and tumor multiplicity (average number of tumors per mouse) were determined weekly until multiplicity reached a plateau (29 weeks). Carcinoma incidence and average carcinomas per mouse were determined weekly from initial detection until 50 weeks after tumor initiation. Squamous cell carcinomas (SCC) were confirmed by histopathology.

#### C. Standard/Delayed/Embryonic Timing of Exposure Experiments

Standard Cohort: FVB/NJ mice were purchased from the JAX labs. The standard model is based on a well-established two-step carcinogenesis models in which adult WT mice are exposed to topical DMBA at 8 weeks of age followed by twice weekly administration of topical TPA for an additional 20 weeks. Mice are subsequently observed for tumor formation and sacrificed once tumor(s) are observed.

Embryonic Cohort: FVB/NJ mice were purchased from the JAX labs. Animals were paired at 8-10 weeks. Females with positive vaginal smears(spermatozoa) are given 4 doses of DMBA (15mg/kg) by oral Gavage at days 14 to 17 of Pregnancy. Litters are divided in two groups the first group at 8 wks. old was treated biweekly promotion with TPA (200 µl of a 10–4 M solution in acetone) for 20 weeks and the second group was monitored for 20 weeks.

Delay cohort: FVB/NJ 8 to 10 weeks old mice were purchased from the JAX labs, first group received a single dose of DMBA (25 μg per mouse in 200 μl acetone applied to shaved dorsal back skin). And waited 25 weeks before commencing treatment with TPA (200 μl of 10−4 M solution in acetone) twice weekly for 20 weeks. second group received a single dose of DMBA (25 μg per mouse in 200 μl acetone applied to shaved dorsal back skin). One week after initiation tumors were promoted with TPA (200 μl of 10−4 M solution in acetone) twice weekly for 20 weeks.

#### D. Cisplatin Treatment

FVB/NJ mice were purchased from the JAX labs. Cisplatin (cis-Diammineplatinum(II), D3371) was purchased from TCI and was dissolved in sterile saline solution at 0.8 mg/ml. A single dose of DMBA was applied to the back skin of the mice at 8 weeks. Mice were then treated with TPA twice a week for 10 weeks followed by tumor formation. Approximately 4 weeks later mice were injected IP with three doses of 100 ul cisplatin or saline solution administered at 4 days interval (day 1, 5 and 9); cisplatin was dosed at 5ul/g of body weight at the time of injection.

#### E. TPA/Wounding Using Inducible Tissue-Specific Activation of Ras mutations

FVB/NJ mice were purchased from the JAX labs. For wounding experiments, 8- to 10-week-old LGR5, Lgr6-EGFP-CreERT2+/−/LSL-KrasG12D, LGR5, Lgr6-EGFP-CreERT2+/−/LSL-HrasG12V mice were administered one topical dose of 4OHT (1 mg in 200 μl of 100% ethanol) (Sigma-Aldrich) on back skin to induce Cre expression, followed by full thickness skin wounding 7 days later. For TPA experiments, skin tumor development 8- to 10-week-old LGR5, Lgr6-EGFP-CreERT2+/−/LSL-KrasG12D, LGR5, Lgr6-EGFP-CreERT2+/−/LSL-Hras mice was initiated with a single dose of 4OHT (1 mg in 200 μl of 100% ethanol) on the dorsal skin at 8 weeks of age, followed by biweekly promotion with TPA (200 µl of a 10–4 M solution in acetone) for 20 weeks. Papilloma number was recorded up to 20 weeks. Mice were sacrificed when carcinomas develop reached a size of >2 cm in longest diameter, papillomas were removed from skin, and all internal tumors were resected.

With the exception of the Dietary cohort, animals were housed in standard conditions in accordance with the rules and protocols stipulated by the UCSF Institutional Animal Care and Use Committee (IACUC) and all mouse experiments were approved by the University of California at San Francisco Laboratory Animal Resource Center. Mice were euthanized when carcinoma reached 2 cm in diameter or meet the humane or experimental end point such as weight loss according to rules and protocols stipulated by UCSF Institutional Animal Care and Use Committee. The Dietary cohort animal protocols were independently reviewed and approved and were housed in standard conditions in accordance with the rules and protocols stipulated by Texas A and M University.

### Tumor Sample Processing and DNA extraction

All tumors were reviewed by a board-certified pathologist. DNA was extracted using Qiagen kits using standard procedures followed by library preparation and quality assurance steps are as described previously(6,63)

### Sequencing

We performed whole genome sequencing of 93 tumor samples with matched normal tail tissue from the same animal for all of the cohorts with the exception of the tumors in the dietary cohort (N=14), in which case 10 normal tail tissue from animals of the same syngeneic background were sequenced as controls. Illumina HiSeq X Ten platform generating 151 base pair paired-end reads. Sequencing reads were aligned to the reference mouse genome (GRCm38) using BWA-MEM(64). Sequence coverage was 31.75-47.79 fold (median 38.78) after duplicate removal.

### Variant calling

The variant calling algorithms of the Cancer Genome Project, Wellcome Sanger Institute, were used with default setting: cgpCaVEMan24 for base substitutions(65); cgpPindel25 for indels(66); and BRASS (https://github.com/cancerit/BRASS) for structural variants. We performed additional post-processing steps to eliminate false positive calls, due to technology specific artefacts and germline variants. For all samples, we removed base-substitutions with a median alignment score of mutation-reporting reads (ASMD) < 140. To perform signature analysis, we filtered out variants present in multiple tumors for samples in the dietary cohort, in order to decrease the likelihood of contaminating SNPs since these samples did not have animal specific matched normal. This filter was not used for the driver gene analysis.

### Extraction of mutational signatures and decomposition to known human mutational signatures

We used SigProfilerMatrixGenerator(24) to categorize mutations into classes and to plot mutational spectra. De novo substitution, doublet/dinucleotide and INDEL signatures were extracted using SigprofilerExtractor (https://github.com/AlexandrovLab/SigProfilerExtractor), which is based on a non-negative matrix factorization method. Before comparing the extracted mouse mutational signature to COSMIC database signatures, we performed a normalization, multiplying signatures by the human mutational opportunity (hg19) and dividing them by the mouse mutational opportunity (mm10). After de novo signature extraction, we used SigprofilerExtractor to decompose these signatures into known human signatures in the COSMIC mutation database.

### Resampling Reduction for Mutation Types

The mutational catalogue of a cancer genome, defined over an alphabet of mutation types Ξ with *k* letters, can be mathematically expressed as a set *S* = {*s*_1_, *s*_2_, …, *s*_*n*_}. To perform a relative reduction of *p* for a given set *Q* of *l* mutation types defined over the same alphabet Ξ, where *Q* is a proper subset of *S*, the number of somatic mutations of each type, *i.e.*, *q*_*i*_ ∈ *Q*, were reduced by applying Poisson resampling with parameters *λ* = *p* ∗ *q*_*i*_. After performing the reduction, samples were subjected to set of routine mutational signature analyses.

### Driver gene analysis

We intersected CaVEMan filtered calls against the homologues of a previously published list of 369 driver genes in human cancers(36). We selected mutations that altered coding sequence: missense, nonsense, splice site mutations, start lost or stop lost substitutions. Custom analyses and visualizations were performed using the *R* library *Oncoplot*.

### Expression analysis

RNA was generated from frozen tumor samples from high and low BMI male animals. Tail tissue was frozen in liquid nitrogen, ground by chilled mortar and pestle, and suspended in TRIzol. TRIzol RNA extraction was then performed, followed by purification by Qiagen kit. RIN values were calculated by Bioanalyzer and samples with high quality (greater than six) were used for analysis.

Samples were hybridized to an Affymetrix mouse gene ST 1.1 array. Gene expression values were calculated in R using the oligo package (1.26.6)(67). Expression values were normalized at the transcript level via the RMA algorithm using a custom set of probe annotations. This custom set was designed to account for the high quantity of sequence dissimilarities between FVB and SPRET mice by removing any probe with a known single nucleotide polymorphism (SNP) between these two strains from the annotation(32). Expression levels were then normalized across plates via the COMBAT method(68).

We used the software Carmen, which has been described previously(32,69), to compare the expression levels of genes independently in males, females, and a combined dataset. Because the phenotype of interest, namely association between obesity and increase risk of carcinogenesis and tumor progression was a strongly sexually dimorphic trait observed in males, we opted to show expression analysis results for the analysis in male mice. We performed this analysis for normal (back skin) and tumor tissue to identify the genes with the strongest expression correlation with high or low BMI.

All other analyses and plotting were performed using *R* utilizing custom scripts.

## Acknowledgements

The authors would like to acknowledge David Adams, Jon Teague and Laura Humphreys as well as other members of the Sanger Institute for discussions, and administrative and technical support. This work was funded by the Mutographs Cancer Research UK Grand Challenge Award (C98/A24032; (PI Dr. Michael Stratton)), the CRUK/NCI PROMINENT Cancer Grand Challenge Award, US National Cancer Institute (NCI) grant R35CA210018, and the Barbara Bass Bakar Professorship of Cancer Genetics (to A.B.). YRL was supported by the NIH/NCI F32 post-doctoral fellowship F32CA232635 and an NIH/NCI Paul Calabresi K12 Career Development Award.

## Author Contributions

The study was conceived and supervised by AB. Tumor models were selected/generated by EK, RD, and the caloric restriction and high fat diet samples were provided by JD. Cisplatin treatment was carried out by DW, RD, and DNA/RNA isolation and gene expression profiling were carried out by KH, MQR and QT. NB and OM provided technical assistance and advice, and computational analysis was performed primarily by YRL, with assistance from LR. Mutational signature analysis was carried out by YRL, MI, and SB, under the supervision of LA. Histopathology evaluation was performed by EK and RD and sequencing was performed at the Sanger institute. the manuscript was written by YRL and AB with contributions from the other authors.

## Data Availability

The raw sequencing data are available for download from the European Nucleotide Archive (https://www.ebi.ac.uk/ena/browser/home) under accession nos. ERP024409, ERP024410, ERP107157, ERP110302, and ERP118253.

## Code availability

The code used in this study is available upon request.

