## Supplemental figure for "The interplay between dormant mutated cells and tumor promotion by chronic tissue damage in determining cancer risk"

**Supplemental Figures and Tables**

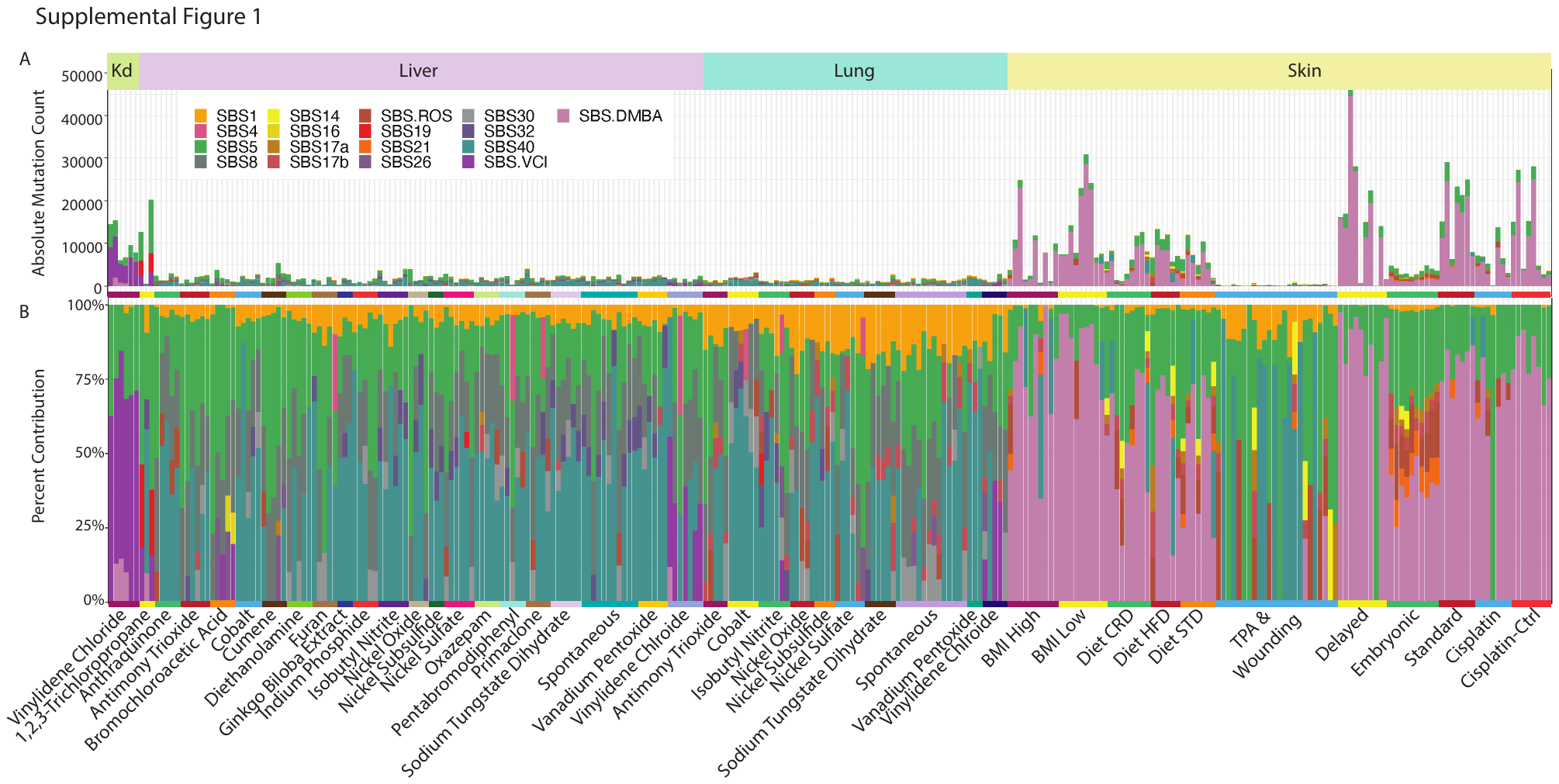

**Supplemental Figure 1 SNV mutations, functional mapping of mutations and mutational signature assignments using the COSMIC mutational signature database.**

Results from SBS mutational signature analysis combining whole genome sequencing data from 188 mouse tumors induced by chronic exposure to suspected environmental carcinogens and the 107 tumors from the present study. Colours denote the contribution of mutations by mutational signature assignment after decomposition of known COSMIC signatures. Each column denotes a sample’s absolute (TOP) and relative (BOTTOM) mutational load.

**Supplemental Figure 2 SBS mutational signature analysis removing 90% T>A mutations.**

A. Absolute SNV mutation load for each tumor (each bar) with colours denoting assignment to each SBS mutational signature after decomposition to known mutational signatures in the COSMIC database.
B. Relative contribution of SNV mutations by COSMIC mutational signature type for each tumor sample. Note the increased prevalence of SBS.ROS once the number of total mutations contributed due to DMBA are reduced.

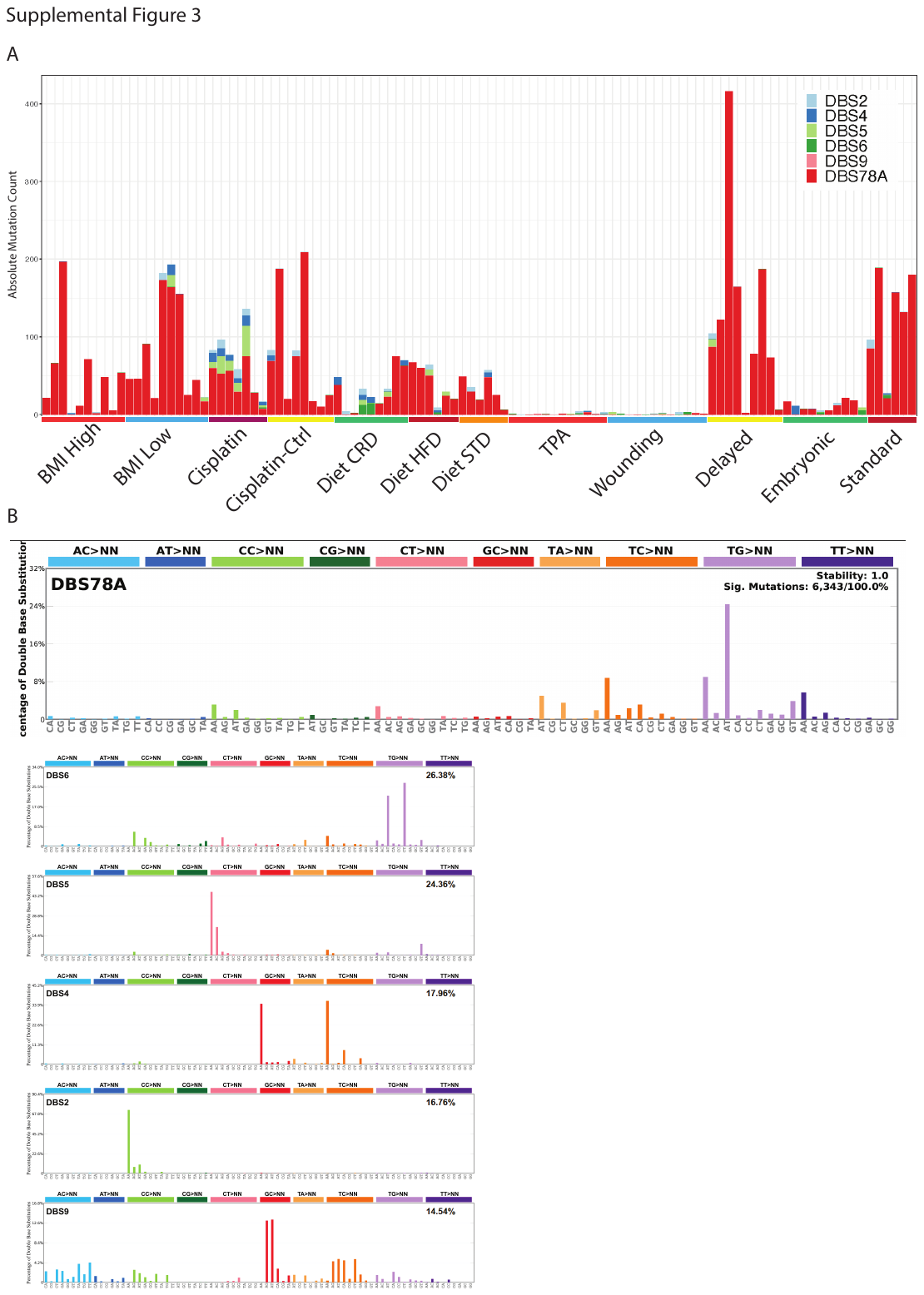

**Supplemental Figure 3 Dinucleotide mutational signature analysis and mutational signatures**

A. Absolute count of dinucleotide mutations coloured by the decomposed signatures with each column denoting one sample.
B. Decomposed dinucleotide mutational signatures identified in the analysis including DBS78A which is a novel dinucleotide signature and DBS5 which is the dinucleotide signature identified as being associated with cisplatin exposure.

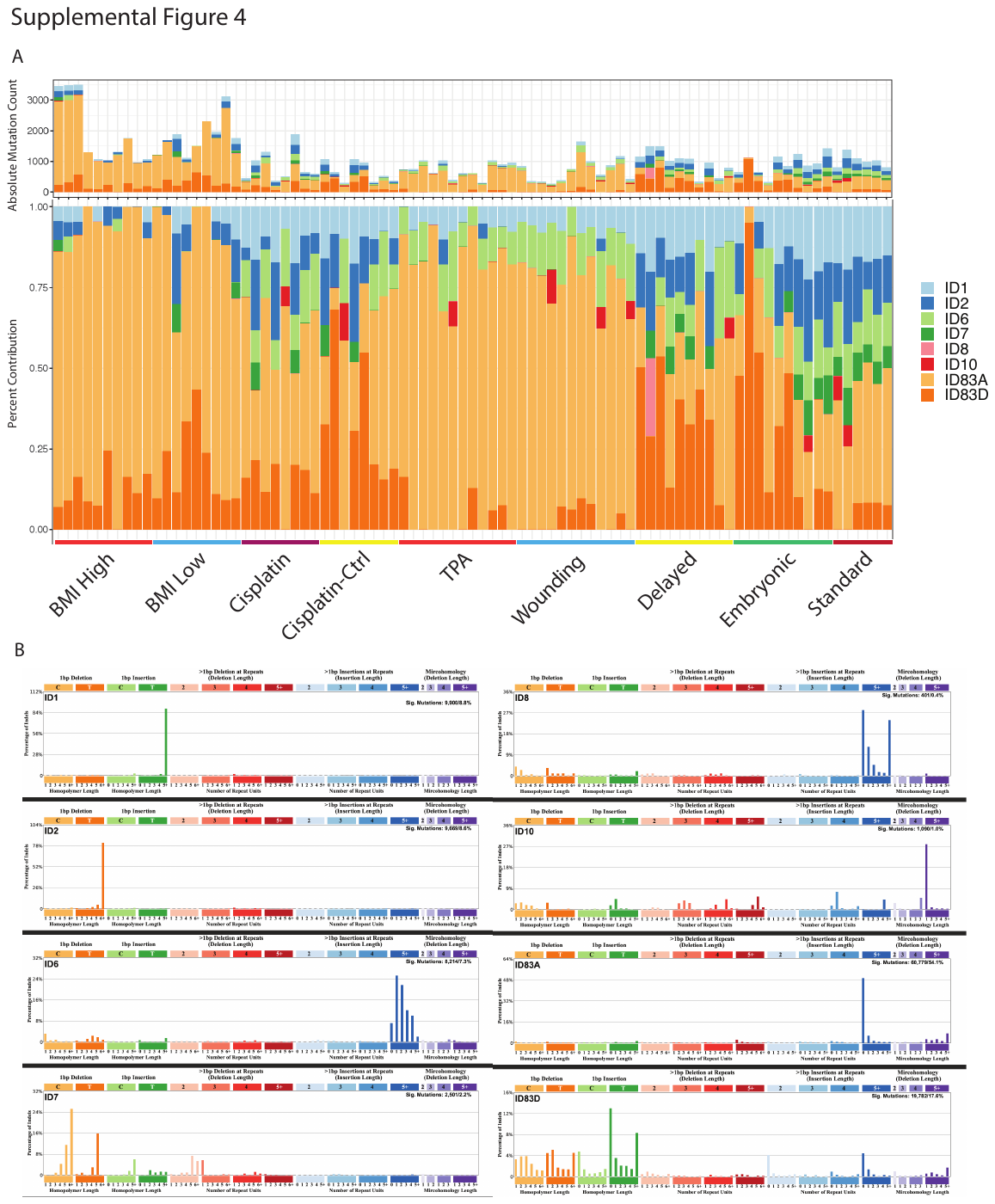

**Supplemental Figure 4 Indel mutational signature analysis and mutational signatures**

A. Absolute (TOP) and relative (BOTTOOM) mutation load coloured by decomposed COSMIC InDel signatures across analysed samples. No specific InDel signature was associated with any of the models examined.
B. Decomposed mutational signatures identified in the analysis.

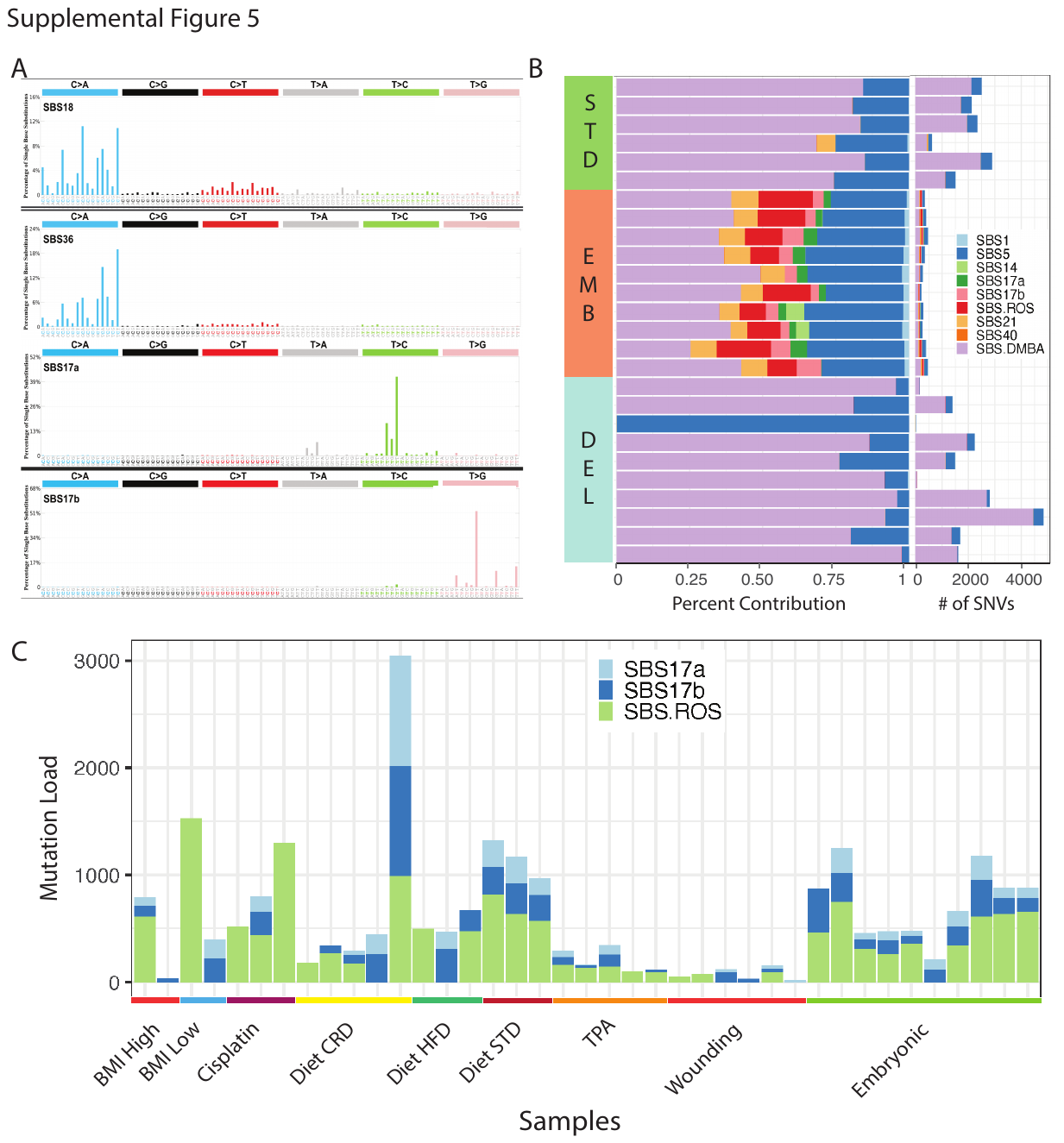

**Supplemental Figure 5 Delay and in-utero exposure models suggest a role for epigenetic modifications in the repair of mutations attributed to ROS during embryogenesis.**

A. COSMIC mutation signatures that have been previously attributed to ROS mediated DNA damage; SBS 18 and 36 are thought to be related to ROS while SBS 17a and 17b were more broadly thought to be related to generalized recurrent inflammation.
B. Significant contribution of SBS/ROS and SBS17a/b mutations to the embryonically derived samples but no other samples in the standard or delayed DMBA/TPA two step skin carcinogenesis models.
C. Contribution of each of the four COSMIC signatures associated with ROS across tumor samples (only samples with at least one mutation associated with SBS17 or SBS.ROS are included).

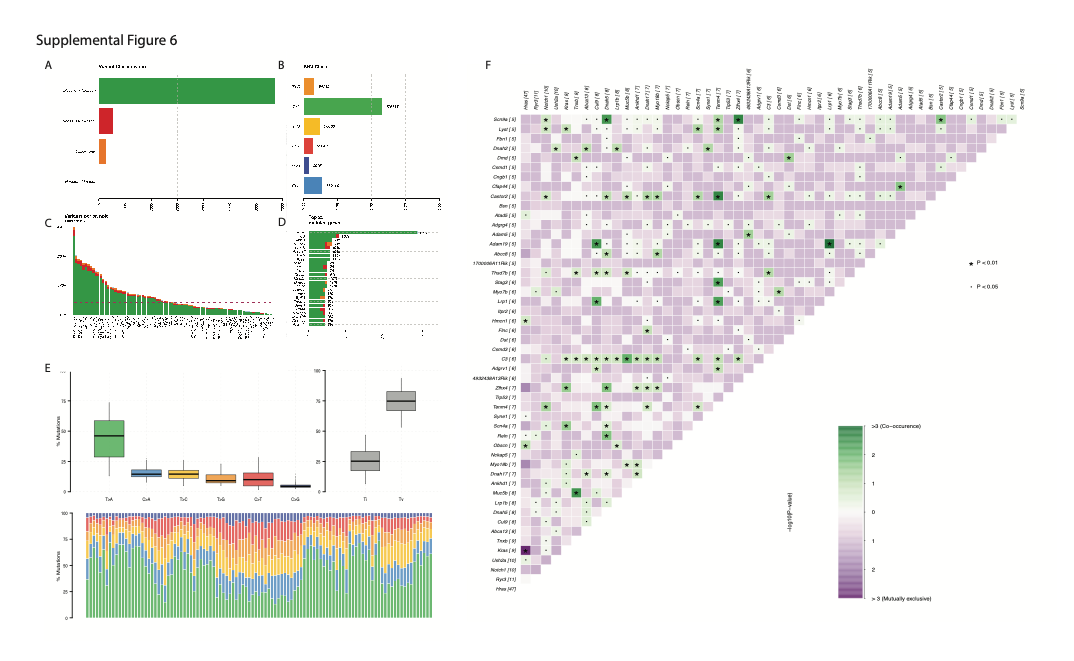

**Supplemental Figure 6 Driver mutation analysis**

A. Classification of mutations after filtering for nonsynonymous mutations mapping to genes previous reported to be cancer driver genes.
B. Driver mutations categorized by nucleotide changes.
C. Bar graph of tumor samples in decreasing order by total number of driver variants per sample and coloured by mutation type.
D. The 25 most frequently mutated genes plotted in decreasing order by the number of unique instances observed in the tumor samples coloured by the type of mutation.
E. Average number of observed mutations by nucleotide change and type (TOP) and per sample distribution of types of nucleotide change observed (BOTTOM).
F. Co-mutation plot of the most commonly mutated genes; colour scale based on whether mutations are mutually co-observed (green) or mutually exclusive (purple). Asterisk (p<0.01) and dot (p<0.05) denote statistically significant.

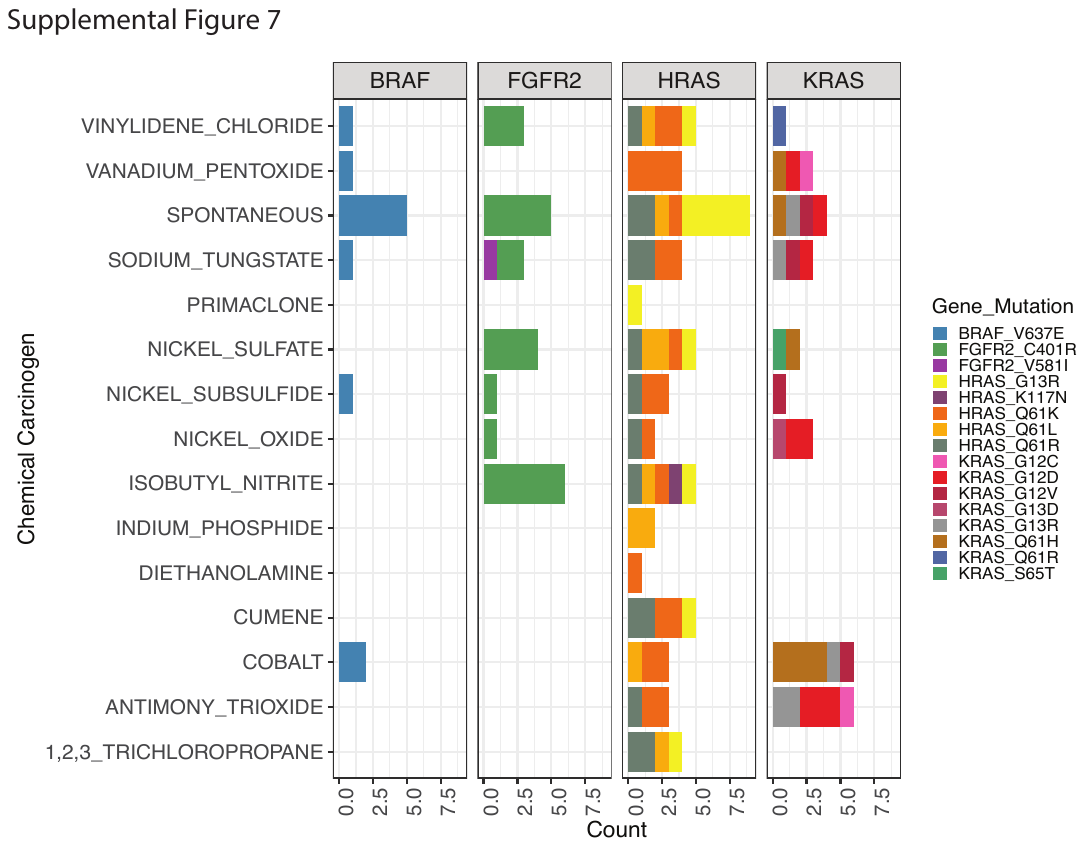

**Supplemental Figure 7 Driver mutations observed in chemically induced tumors or spontaneously derived tumors**

Recurrently observed mutations in the four most common driver genes across the 188 chemically induced or spontaneously arising tumors (liver and lung) coloured by the driver mutation and specific amino acid context change. See methods as described in and data accompanying Riva et al. (6)

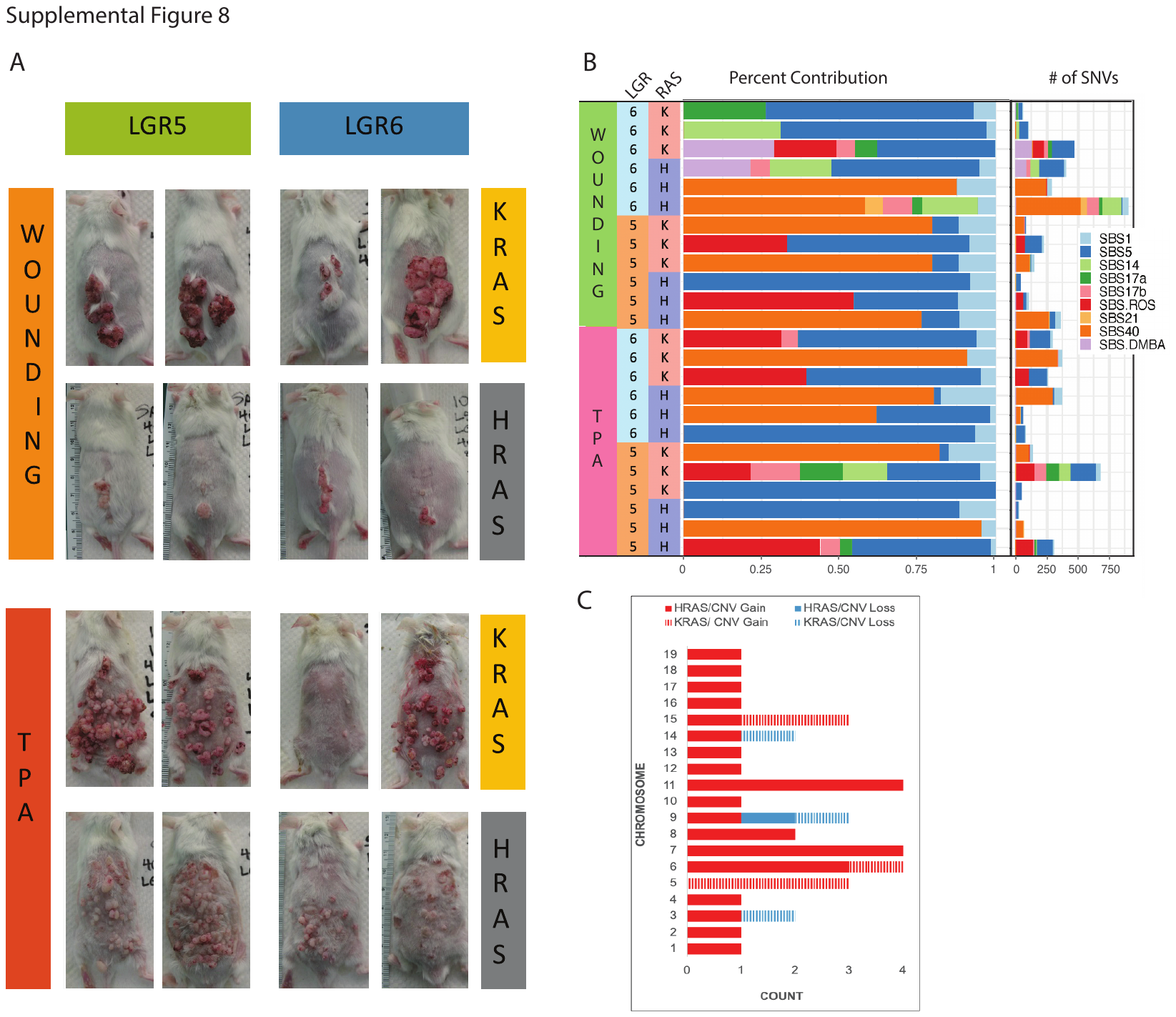

**Supplemental Figure 8 Inflammation is the time limiting step in tumorigenesis in LGR5/6-Cre activated KRAS/HRAS mouse.**

A. Representative tumors induced using genetic models in the absence of chemical mutagens. Animals are heterozygous for LGR5 or LGR6 driven expression of either HRAS or KRAS. More tumors are observed typically in KRAS mutated animals. Tumors observed in wounding animals are generally larger while many smaller tumors are more typically observed in animals exposed to TPA.
B. Absolute (Right) and Relative (Left) SBS decomposed Cosmic mutational signature analysis.
C. Large CNVs/Aneuploidies identified across samples by chromosome and coloured based on driver mutation (KRAS vs HRAS) and a gain or loss was observed.

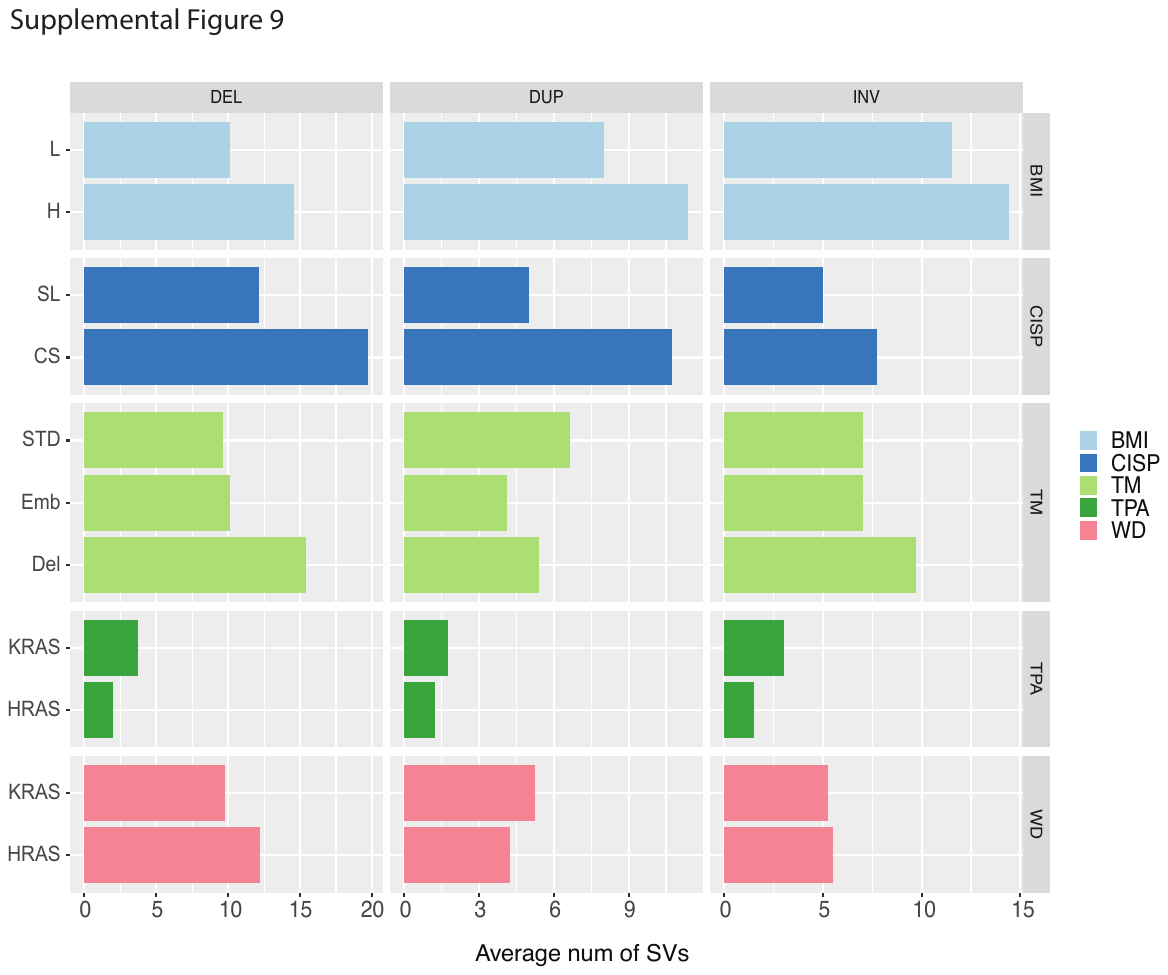

**Supplemental Figure 9 Structural variants identified by WGS.**

Average number of structural variants identified by type. Dietary samples were removed from analysis due to lack of matched controls for each individual tumor.

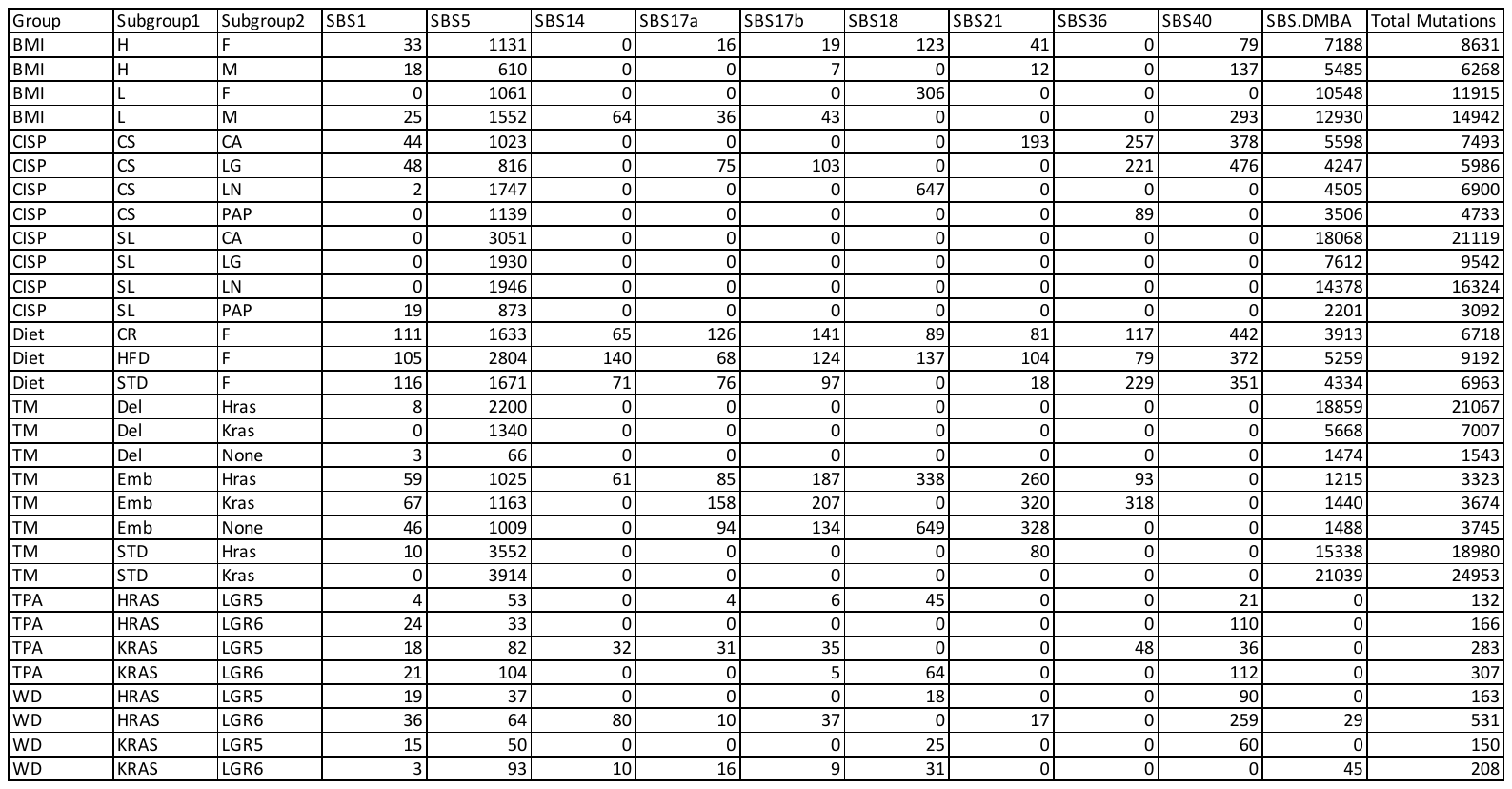

**Supplemental Table 1**

Breakdown of average mutations per tumor assigned to each decomposed SBS signature based on sample groupings.

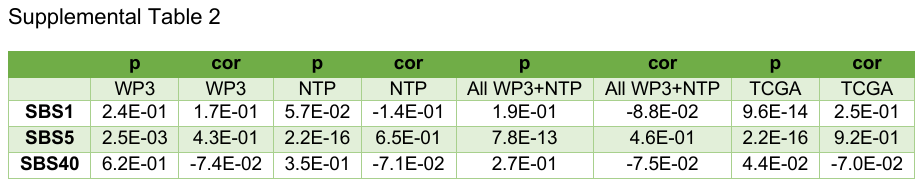

**Supplemental Table 2**

Comparison of association between total mutational load and each of the “clock” or “clock-like” mutational signatures in the WP3 (present cohort), NTP+WP3 (all mouse tumors including those from chronic exposure to carcinogens), and TCGA. Only SBS5 appears to be strongly associated with total mutational load in all instances.
